# Human vicarious trial and error is predictive of spatial navigation performance

**DOI:** 10.1101/332189

**Authors:** Diogo Santos-Pata, Paul FMJ Verschure

## Abstract

When learning new environments, rats often pause at decision points and look back and forth over their possible trajectories as if they were imagining the future outcome of their actions, a behavior termed “Vicarious trial and error” (VTE). As the animal learns the environmental configuration, rats change from deliberative to habitual behavior, and VTE tends to disappear, suggesting a functional relevance in the early stages of learning. Despite the extensive research on spatial navigation, learning and VTE in the rat model, fewer studies have focused on humans. Here, we tested whether head-scanning behaviors that humans typically exhibit during spatial navigation are as predictive of spatial learning as in the rat. Subjects performed a goal-oriented virtual navigation task in a symmetric environment. Spatial learning was assessed through the analysis of trajectories, timings, and head orientations, under habitual and deliberative spatial navigation conditions. As expected, we found that trajectory length and duration decreased with the trial number, implying that subjects learned the spatial configuration of the environment over trials. Interestingly, IdPhi (a standard metric of VTE) also decreased with the trial number, suggesting that humans benefit from the same head-orientation scanning behavior as rats at spatial decision-points. Moreover, IdPhi captured exclusively at the first decision-point of each trial, was correlated with trial trajectory duration and length. Our findings demonstrate that in VTE is a signature of the stage of spatial learning in humans, and can be used to predict performance in navigation tasks with high accuracy.

## Introduction

During spatial navigation experiments, rats often stop at the maze intersections to look back and forth toward their possible route choices. Such behavior, originally interpreted as an exploration of possible future outcomes, was termed ‘vicarious trial and error’ (VTE)^1,2^. It has been shown that animals exhibit more VTE behaviors during early phases of spatial exposure^2^, and when discriminating between possible options becomes more difficult^1^.

From a spatial navigation perspective, VTE has been linked to the firing of hippocampal place cells. Place-cells are neurons in the hippocampal formation whose receptive-fields are tuned to specific locations of the explored environment^3^. Such position-related signals have been shown to be used when rats probe future trajectories during spatial-decision making processes^4^. Specifically, during VTE behavior occurring at navigational intersections, hippocampal place-cells are sequentially activated in the direction of the maze arm towards which the animal orients itself. Such findings gave rise to the hypothesis that VTE serves the purpose of setting the neuronal mechanisms of spatial representation for simulating future trajectories rather than simply potentiating the acquisition of sensory information.

Rodent spatial learning has been described to employ two types of navigational strategies^5^: “place” and “response”. “Place” strategies use environmental sensory information to encode current and future locations and leads to the ability to adapt to environmental sensory changes. “Response” strategies, on the other hand, rely on the learned sequence of motor-actions to reach a goal-location in a given navigational context. It has been shown that the different stages of spatial learning recruit one or the other strategy, with each using its own distinct, associated neural mechanism (see^6^, for a review on VTE and spatial learning). Specifically, lidocaine inactivation of hippocampal neural populations impaired “place” strategies and striatal caudate nucleus inactivation disrupted “response” navigational strategies^7^. The distinct recruitment of these brain regions during spatial navigation suggests that with increased familiarity, initial learning stages, when tasks still require the association of spatial and sensorial information for the understanding of the experimental contingency, progressively transition into a cue-action coding scheme^5^.

Situating oneself in space is dependent on one’s exploration history and perception of environmental cues. As navigational learning increases, attention allocated for sensory precessing decreases and motor-outcome memories play a more prominent role^6,8^. However, the signatures of such learning/relearning and how experience affects them remain unclear. Head-scanning behaviors have been shown to relate to novelty detection and are suggested to serve the purpose of sensory acquisition^9^. Even though VTE has been mostly studied in rodents’ spatial navigation, humans might also benefit from, and exhibit, such behavior. At the very least, the neuronal apparatus serving spatial representation has been found in the human hippocampus^10^. Also, at the behavioral level, human saccade–fixate–saccade sequences have been reported during moments of decision-making in visuospatial tasks, suggesting that humans also project onto their possible options before taking a decision^11,12^ Thus, one could expect that similar to rodents, specific behavioral mechanisms take place during the various stages of spatial learning in the human model.

To test whether humans take advantage of VTE behaviors during spatial navigation, we devise a virtual reality navigational task where participants are asked to perform goal-oriented navigation. We test their ability to perform goal-oriented navigation within a virtual environment where exposure conditions promoted either “place” (through low-frequency exposure to a navigational contingency) or “response” (through high-frequency exposure instead) strategies. We hypothesize that head-orientation during spatial decision-making is a behavioral correlate of learning and, moreover, that it is predictive of navigational performance.

## Methods

### Experimental design

Twenty subjects (8 female, age 27 ± 3 years old, right-handed) were recruited from Universitat Pompeu Fabra student’s community. The experiment consisted of 62 goal-oriented virtual navigation trials, where subjects were asked to perform a spatial memory task by reaching a goal-object location the quickest possible. A squared virtual environment (20 virtual meters (vm) on each side), made up of corridors and solid walls, was built using the Unity3D game engine (San Francisco, California). The layout was based on the one presented in^13^: a highly symmetrical environment with sparse visual landmarks (fig:1-A), and multiple decision-making points between start- and target-locations. In order to perform virtual navigation, participants controlled a virtual character in first-person perspective. Translation and rotation in virtual space were controlled with a keyboard, with arrow keys indicating the direction of both translation (front/back) and rotation (left/right). Head orientation was controlled with an optical mouse along the physical horizontal plane and had a consistent mapping rate to the virtual world. Head and body rotation were programmed with specific rates of angular changes. Body rotation was set at 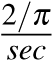of key pressing, while the head angular change was set at 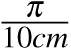 of the optical mouse translation. Therefore, performing virtual direction-scanning behaviors was faster by simply manipulating the character’s head rather than its body. Indeed, such manipulation served the purpose of promoting head rotation when participants sought to acquire environmental sensory information during spatial decision-making. A 1 vm length virtual pole projecting ahead of the users viewpoint was included so that participants were aware of both the body and head orientation in the environment. The virtual speed of navigation was kept fixed throughout the experiment. Visual cues, composed of circular patches of different colors, were uniformly distributed across the virtual maze and were the only position-relevant information available (see a screenshot of the virtual maze in figure:1-B). Two possible starting locations, as well as two distinct target locations, were defined. Therefore, there were four possible start-target combinations, all of which were tested. Because rodent VTE has been linked to spatial learning and consequently to place/response navigational strategies, we assigned each of the four start-target combinations to different exposure frequencies. Subjects were exposed to the high-frequency condition in 60% of the trials and to the low-frequency condition in 28% of them. The remaining two start-target combinations were interleaved with the high- and low-frequency conditions and were not considered for further analyses. Trial types were uniformly shuffled across the experimental session and could not be predicted by the subjects. At every trial, a visual icon representing the target object (color and shape) was presented and kept visible on the computer screen (fig:1-B). Each trial started with the virtual character placed at one of the two possible starting locations and ended when the goal-object was reached. At the end of each trial, a blank window with a white cross centered on the screen was displayed for 4 seconds. After that, the white cross disappeared, indicating that participants could now start a new trial by pressing any of the arrow keys on the keyboard.

**Figure 1.**
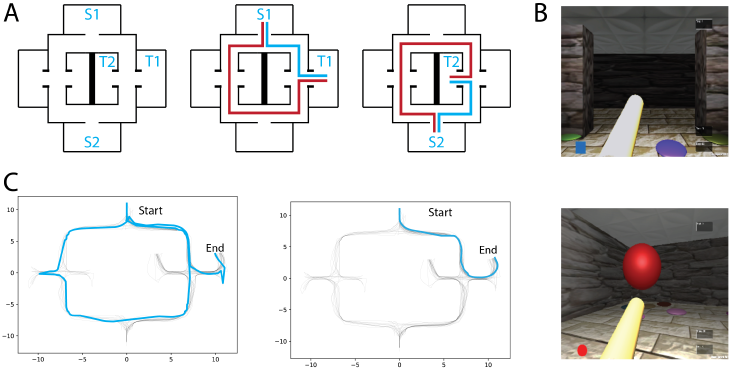
**A** Left: Virtual maze layout. S1,S2 and T1,T2 denote the possible starting and target locations, respectively. Two exmaple of starting-target combinations with respectively optimal (blue) and erratic (red) trajectories schematics. **B** Screenshots of the environment through first-person perspective. **C** Example of an early (left) and a late (right) trial trajectories from one subject.

### Navigational measures

#### 0.0.1 Trial duration and distance

The position and angular orientation of the virtual character were recorded throughout the experiment at a sampling rate of 70HZ. Trial duration was considered as the elapsed time from the trial onset till the moment the character reached the target-object. The trial distance was computed by summing the traveled distance given by:

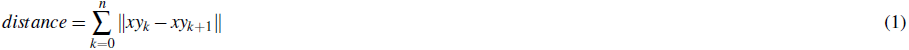

where, *xy* are the spatial coordinates of each point *k* in the trajectory.

#### 0.0.2 Route difference

The route difference, a measure of similarity between pairs of navigational trajectories was used to quantify learning and automation. Each trajectory consisted of the path taken between the starting location and target location of each trial. Therefore, the route difference reflects the Euclidean distance between the two given navigational trajectories. In order compute the route difference between a trajectory pair, each trajectory was clipped to the vector length of the shortest trajectory and interpolated along each data-point. Thus, their difference was measured between each aligned data point for every pair of trials. The final score was obtained by averaging the difference between trials i and j given by:

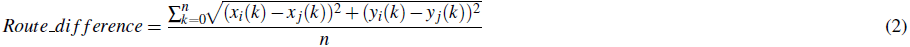

where, *x* and *y* are the virtual environment x-axis and y-axis coordinates of each point *k* from either *i* or *j* trajectories, and *n* is the total amount of positions from the interpolated trajectories.

#### 0.0.3 IdPhi: VTE behavior

The IdPhi measure has been used in the rodent literature to quantify VTE behaviors at decision-points of navigational tasks^5^. We adapted IdPhi to our dataset in order to quantify changes in angular orientation at decision-points during navigation. Navigational components used in the IdPhi analysis comprised the trajectory segments at the moment the virtual character approached the first spatial decision-point until it entered one of the arms of the maze. The first derivative of the character’s head orientation was extracted to obtain changes in angular orientation 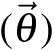 and unwrapped to avoid circular transitions. The IdPhi measure was then calculated by integrating the absolute values of |*dPhi*|, given by:

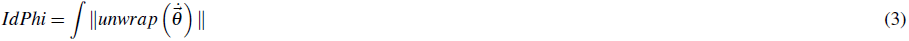

where *θ* are the virtual character’s angular orientation points of the trajectory segment included in each analysis.

#### 0.0.4 Head-orientation variability

The IdPhi measure of VTE behavior extracts a single value for a given trajectory segment. In order to observe how participants modulated their head-scanning behavior along with their trajectories, we measured changes in head-orientation and quantified its modulation in their spatial rate dimension for each trajectory. We first applied a band-passed filter (1-20Hz) onto the head-orientation signal. The resulting orientation signal was z-scored and its envelope was extracted via the Hilbert transform. The orientation amplitude was normalized to its corresponding trial maximum amplitude. The environment was segmented into 30*x*30 virtual bins (bin size = 1 vm) and the orientation amplitude score was normalized by the time spent at each of the environment’s spatial bins. The resulting orientation variability rate maps were then smoothed using a 2D Gaussian filter (*σ* = 1.2).

## Results

### 0.1 Spatial learning

All analyses and data processing were done using the scipy^14^ and numpy^15^ python packages. Spatial learning was measured through trajectory similarity across trials of the same (high/low frequency) condition. For illustrative purposes, two trajectories from the same participant, one from an early trial and one from a late trial, are depicted in figure:1-C. As expected, early trials of both conditions tend to take longer to complete than later trials in the same condition, suggesting that overall, participants were capable of understanding the environment’s layout and task contingencies (fig:2-B). Navigational duration plateaued after the first 6 trials (23.1 ± 3.6 sec for the low-frequency condition, and 15.29 ± 1.35 sec for the high-frequency condition). Similarly, there was little change in trajectory length after the first 6 trials (34.01 ± 4.62 vm for the low-frequency condition, and 24 ± 742.08 vm for the high-frequency condition). Despite these decreases in duration and length, we expected that trajectories would become more similar as participants learned the environmental contingencies (for an example of consecutive high-frequency condition trials from one subject, see fig:2-A). We have measured the route difference between every trajectory pair from both conditions per subject (fig:2-C). Overall, early trials exhibited a greater route difference (low condition *max* = 7.18*vm*, high condition *max* = 5.96*vm*) whereas later trials were found to be more similar (low condition *min* = 0.47*vm*, high condition *min* = 0.21*vm*). In summary, decreases in navigational paths duration and length, accompanied by increases in trajectory similarity suggests that participants were capable of progressively acquiring knowledge of both the environmental configuration as well as the experimental configuration, adapting their spatial routes and decision-making depending on the start-goal contingency.

**Figure 2.**
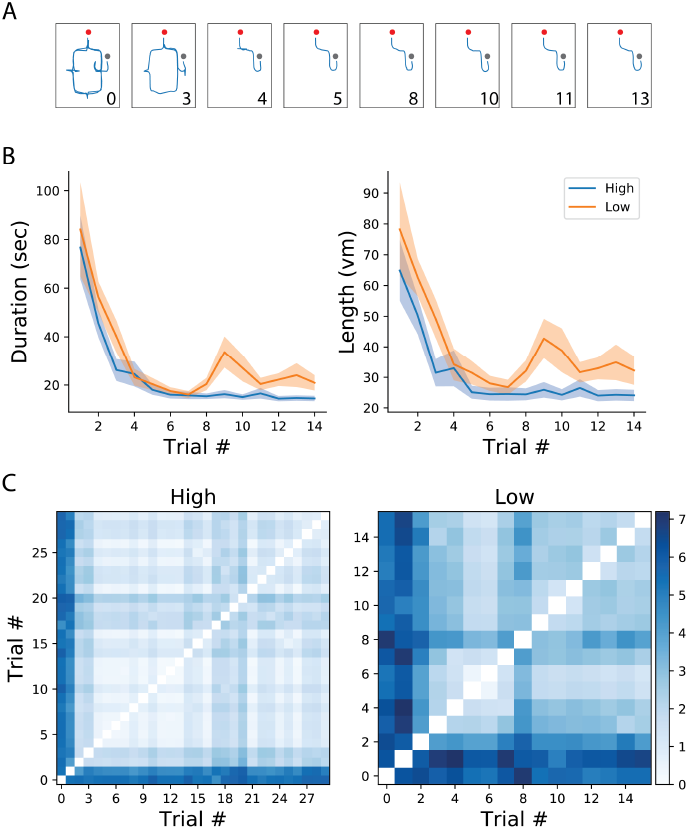
**A** Example of trajectories from one participant in the high frequency condition. Numbers correspond to trial number. Red and gray dots represent the starting and end of the trial, respectively. **B** Mean and standard error of trial duration (left) and length (right) for the first 14 trials of each condition (high/low). **C** Route difference between every trajectory pair in each condition, averaged across subjects.

**Figure 3.**
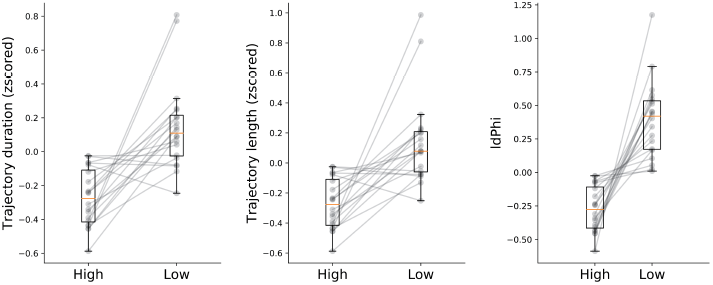
Navigational measures at high and low frequency conditions. Trajectory duration, length and IdPhi were smaller for the high condition when compared to the low frequency condition (t-test: duration *p <* 0.05, length *p <* 0.05, IdPhi *p <* 0.01). Grey lines represent the mean of each subject in a given condition.

### 0.2 VTE

We hypothesized that humans benefit from behavioral correlates of spatial decision-making, specifically from VTE behaviors, as has been previously shown in rodents^1,2,5,16^ In order to measure whether humans do benefit from these back and forth head-scanning behaviors, we quantified navigational variables in both high and low-frequency conditions. Because high frequency conditions have, by definition, a greater probability of occurring, we expected that trajectory length and duration of trials with high-frequency start-target combinations would be shorter when compared with low-frequency conditions. The rationale was that because participants were exposed to the high condition more often than to the low condition, associations between environmental sensory information and goal-locations with their respective repertoire of motor actions would be encoded earlier in the experimental session. Surprisingly, the plateau of duration and length was reached within a similar number of trials for both conditions (fig:2). Despite coherent learning stabilization in the number of trials exposed in both conditions, we still expected trajectory length and duration of navigation to differ across conditions. Indeed, duration was significantly lower (t-test, *p <* 0.05) for the high as opposed to the low frequency condition (High: *mean* = −0.27 ± 0.16*std*, low: *mean* = 0.15 ± 0.25*std*). Moreover, trajectory length was significantly shorter (t-test, *p <* 0.05) for the high condition (*mean* = −0.26 ± 0.15*std*) when compared to the low condition (*mean* = 0.14 ± 0.29*std*). Taken together, these results suggests that higher exposure to the navigational task leads to greater route optimization as compared to fewer exposures.

Other than trajectory duration and length, we were also interested in head-scanning events at decision-points. As described in the methods section, we extracted the trajectory segments when participants were crossing through the first decision point. IdPhi scores were split in high and low frequency conditions (fig:2-right subplot). Interestingly, low frequency trajectories revealed a greater variation in head-orientation at the first decision-point (*mean* = 0.39 ± 0.28*std*) when compared with the high condition (*mean* = −0.27 ± 0.17*std*) at a statistically significant level (t-test, *p <* 0.01).

### 0.3 VTE decreases with learning

We have reported on three measures of spatial navigation: trajectory duration, length, and IdPhi at the first decision intersection. We hypothesized that IdPhi scores were predictive of the participant’s ability to perform a goal-oriented navigational task in a static environment. As expected, we have observed greater trajectory duration, length and IdPhi for the low-frequency condition when compared to the high-frequency condition, suggesting that frequency of exposure modulates the internal representation of the environment and/or experimental contingencies. Head-scanning behaviors have been observed to increase the activity of hippocampal place-cells leading to the potentiation of their receptive-fields at a given spatial location^9^. These results suggest one-trial encoding of experiences mediated by place-cell activity. Therefore, one could expect that head-scanning behaviors serve the purpose of acquiring sensory information in moments when hippocampal spatial representation does not properly, or sufficiently, predict the sensory statistics of the environment. However, whether head-scanning is an all-or-nothing type of behavior or if it follows the progressive modulation, as observed for trajectory duration and length, is still unclear. We calculated averaged scores for the three navigational measures at a trial-by-trial basis for each condition (fig:4). As observed in figure:2, trajectory duration and length quickly decreased with contingency exposure. Again, high-frequency condition obtained a greater trajectory score after reaching its plateau when compared with low frequency trials (fig:4, top and middle). Notably, the IdPhi measure of head-scanning followed a similar trend as the previous navigational measures. That is, it progressively decreased with increasing contingency exposure and scored lower for the high-frequency condition after reaching a plateau compared to the low-frequency condition (fig:4-bottom).

**Figure 4.**
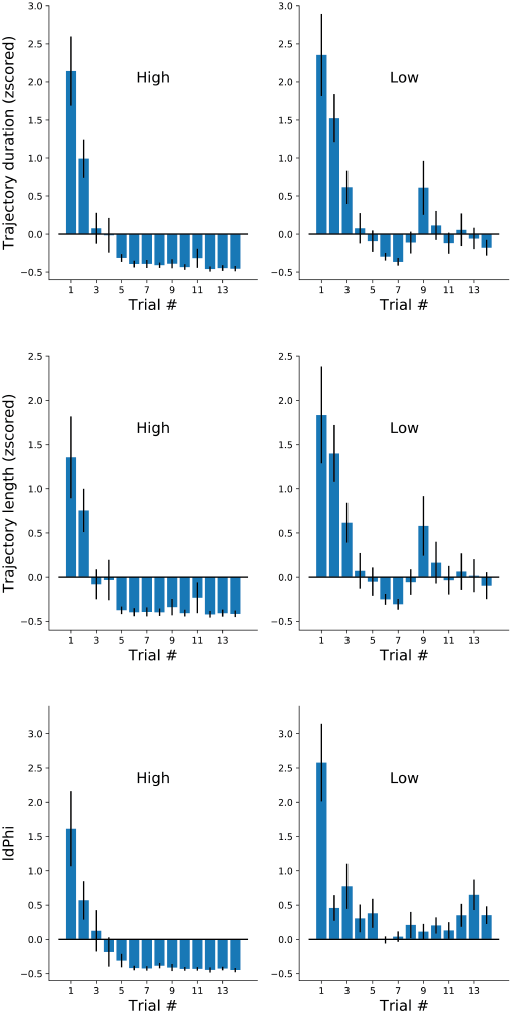
Navigational measures along the first 14 trials of each participant at each frequency conditions. Trajectory duration (mean±sem). Duration, length and IdPhi strongly decreased on the first three trials of each condition. High frequency trials were maintained on negative standard deviation throughout the experiment, while low frequency trials oscillated around zero.

### 0.4 Head-orientation variability is specific to decision-making

GivGiven that in our task the decision taken at the first intersection was of greater importance than at other junctions (optimizing navigational score is heavily dependent on the decision made at the first intersection), VTE was measured by obtaining trajectory segments crossing the first decision point of each trial. In addition, we measured the head-orientation variability within the entire environment (see methods). Put briefly, we computed the first derivative of each trial’s head-orientation in polar coordinates, then applied a low-pass filter and a Hilbert transform on the signal. The amplitude of the head-orientation variability was normalized to the entire trial signal. Rate maps of each trial were built upon the discretized environment (*bin* = 1*vm*) and normalized to the occupancy duration spent in each spatial bin. Rate maps were aligned (rotated) so that the first-intersection position would match across conditions and averaged head-orientation variability across subjects were used for further analysis (fig:5). In sum, low-frequency conditions have a higher rate of orientation variability at the first decision point when compared with the high-frequency condition (fig:5-top). Next, we built a library of rate maps and performed the clustering permutation statistical test. This way, we could assess whether levels of head-orientation at decision points were significantly higher for the low-frequency condition (p-val threshold was set to 0.05). As expected, a significant cluster survived permutation testing in bins surrounding the first decision intersection (fig:5-bottom, dashed pink inset). Similar to our other navigational measures, the amount of time spent at the first decision-making intersection progressively decreased with learning (fig:5-C), which served as a second control for the between-conditions differences in head-orientation variability observed before. Together, these results demonstrate that head-scanning behaviors are functionally engaged in the spatial decision-making as observed in the rodent literature. Moreover, the fact that the cluster center of mass is specific to a single location of the environment and head-orientation is not modulated throughout the environment supports the hypothesis that VTE is part of a deliberation process.

**Figure 5.**
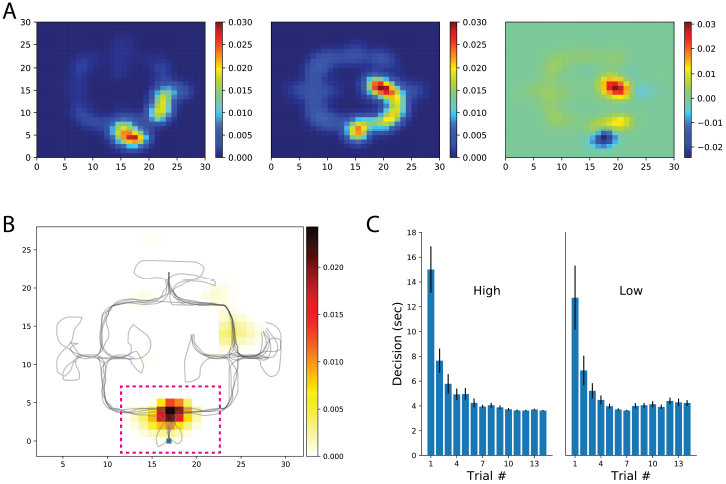
Head-orientation and decision-making duration. **A** Participant’s rate maps were aligned (rotated 180 deg) when necessary so that starting location would match across participants and conditions. Rate maps of low (left) and high (center) frequency conditions, as well as their difference (right) are shown. Shown rate maps represent the mean across participants. **B** Cluster-level statistical permutation test for head-orientation in the low and high conditions. Colored bins are locations that survived multiple comparisons correction with p-value < 0.05. Note the strong cluster at the first decision-point (dashed pink line, zoomed on the right) suggesting a greater head-orientation variance for the low frequency condition. **C** Time spent from trial onset until entrance in one of the possible maze alleys (averaged across participants).

### 0.5 VTE is predictive of navigational performance

We have observed a congruent modulation among the selected navigational measures when comparing between conditions and between trials. However, scoring of trajectory length and duration differ methodologically from the IdPhi scoring in the sense that IdPhi is obtained by a considerably short trajectory segment at the beginning of the navigational path, whereas trajectory length and duration summarize the entire starting point to goal navigational path. Moreover, IdPhi scores progressively decreased on a trial-by-trial basis, suggesting a continuous modulation congruent with spatial learning (fig:4). We have asked whether the amount of head-scanning behavior at the first intersection could be predictive of subsequent navigational performance. As the speed of navigation was programmed to be fixed — a continuous navigation — a small variance in the amount/duration of pauses across trials would have a strong, confounding correlation with trajectories’ length (space) and duration (time). In order to control for navigational continuity, we computed the linear relationship between trajectories’ length and duration through pairwise correlation (Pearson correlation coefficient *r* = 0.96, *p <* 0.01). All participants’ data was z-scored and merged for this analysis). Participant-specific length and duration relationship is depicted in fig:6 top-right. Next, we measured the linear relationship between IdPhi scores at the first intersection with the length and duration of that trial. IdPhi scores revealed to be significantly correlated for trajectory duration (Pearson correlation coefficient *r* = 0.502, *p <* 0.01) and trajectory length (Pearson correlation coefficient *r* = 0.431, *p <* 0.01). Thus, this suggests that head-scanning behavior at early moments of the trial is predictive of the final trajectory performance (fig:6-middle and bottom panels). A measure of group effect was performed by averaging correlation coefficients. Regarding trajectory duration, IdPhi was predictive in 18 out of the 20 participants (*p <* 0.05, *mean* = 0.009 ± 0.005*sem*). Similarly, IdPhi correlated with trajectory length in 17 out of the 20 subjects (*p<* 0.05, *mean* = 0.04 ± 0.02*sem*).

**Figure 6.**
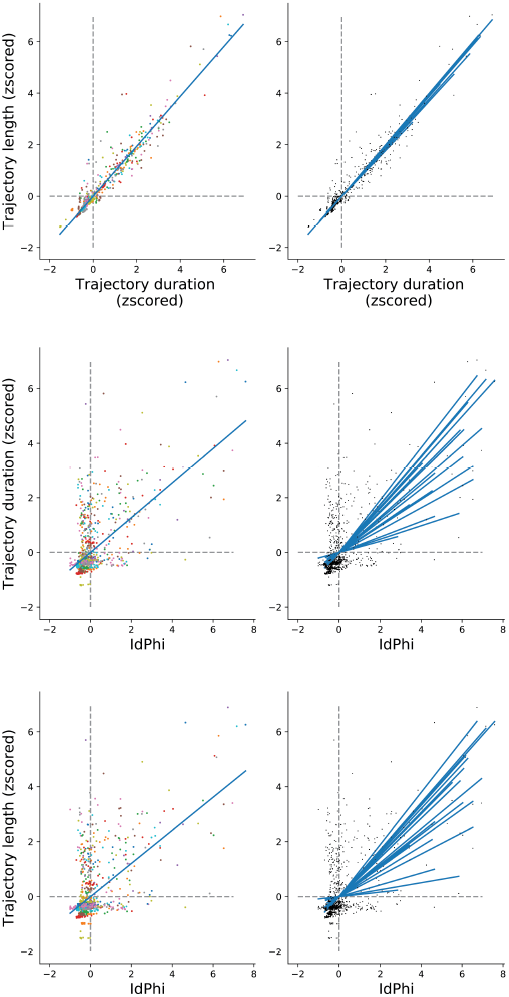
IdPhi at first decision-point correlates with navigational aspects Top. Relation between length and duration of each trial for all subjects (left) is highly correlated (Spearman test *r* = 0.942, *p* = 0). Correlations for individuals are shown on the right column. **Middle** Correlation between length and IdPhi (Spearman test *r* = 0.431, *p <* 0.001). **Bottom** Correlation between duration and IdPhi (Spearman test *r* = 0.5015, *p <* 0.001).

## Discussion

We have tested human spatial learning in a goal-oriented virtual navigation task. Because the behavioral and physiological correlates of animal spatial navigation have been extensively studied in the rodent model, we aimed to quantifiably link the observed rodent behaviors of specifically place/response strategies, VTE and head-scanning behaviors with how humans navigate in space.

Rodent spatial learning has been revealed to be punctuated with very specific behavioral and neuronal markers at distinct learning phases of spatial exploration. One such behavioral marker, the VTE behavior, i.e. pausing at a maze intersection during spatial navigation to look back and forth at the possible options before taking a decision, has been observed for many years^1,2^. Presumably, scanning the environmental configuration while in the process of decision-making suggested that animals were gathering sensory information in order to enrich their world model. Nevertheless, it is known that both rodents and humans are capable of developing neuronal spatial representations of their surroundings^3,10^. Moreover, it has been shown that hippocampal place-cells representing possible future trajectories are sequentially activated during VTE behavior^4^. Therefore, this shows that VTE events occur even when the internal representation of space has been learned, rejecting the hypothesis that VTE serves the sole purpose of boosting the learning of the environmental sensory-statistics at the neuronal level. The hippocampal neural mechanisms for simulating spatial trajectories have been proposed both theoretically and computationally^17,18^ Additionally, a model of spatial representation and trajectory simulation has been implemented in a robotic simulation of the T-maze navigation task and suggested the emergence of VTE behaviors at early learning phases^19^.

Conceptually, pausing at spatial decision-points can be a signature of at least two distinct processes. On one hand, scanning the possible routes to take allows acquiring sensory information regarding the environmental options. This increase in available sensory information could potentially increase the number of hippocampal place-cells tuned to the spatial locations of each of the maze-arms, as observed in^9^, thereby optimizing performance. On the other hand, aligning one’s body towards possible decision options would allow the setting of path-integration hippocampal mechanisms to simulate future trajectories, as has been theoretically and computationally postulated^17,18^. Indeed, hippocampal mental time-travel has been observed to occur when the animal performs VTE behaviors^4^. Regardless of whether VTE behaviors serve to gather spatially relevant sensory information or to modulate internal mechanisms of spatial representation, the fact that they occur at specific spatial learning stages is indicative of their participation in navigational performance^6^.

In order to assess spatial learning, we have devised a navigation task in a highly symmetric virtual environment where the floor, walls and ceiling texture, as well the environment layout would not be sufficient to decode a participant’s avatar’s position. The only reliable sensory signal indicative of position were the visual cues made of circular patches sparsely scattered in the environment that were kept constant throughout the experiment. Because in each trial, participants would start at one of two possible locations and were asked to reach one of two possible targets, the trajectory taken to reach the goal was specific to each of the combinations. The ability to form a spatial representation of the environment and task contingency was measured through spatio-temporal characteristics of the participants spatial trajectories. We observed that participants were capable of improving their trajectories length and duration after a few trials of exposure to a given starting-goal combination (fig:2-B). Similarly, we have observed that with learning, the similarity of trajectories belonging to the same condition tended to increase (fig:2-B,C). Despite the navigational improvement throughout the experiment, such spatial measures do not entirely capture the underlying mechanisms by which participants learned the task. On the one hand, it could be that with exposure, participants incrementally acquired spatial knowledge and were capable of forming a proper spatial representation of their surroundings, therefore becoming capable of situating themselves within the maze, understanding the environment’s layout and remembering spatial locations associated with target objects. On the other hand, such navigational improvements could be due to the participants’ ability to associate a set of landmarks/cues at the initial trial’s location with a sequential chain of navigational actions leading to the target-object.

An alternative hypothesis would be that participants recruit distinct navigational strategies (place/response) accordingly with the trial condition that they are requested to perform. Similar to the alternation of navigational strategies strategies observed in rodents^5^, the levels of IdPhi scoring in our experiment were significantly higher for the high-frequency condition when compared to the low-frequency condition. Therefore, it not only suggests that participants adapt their strategies to the experimental contingency but also rejects the hypothesis that participants relied on a memory of action sequences in the low-frequency condition. Our results are congruent with the assumption that humans take advantage of head-orientation during spatial-decision making, a component observed in both VTE and head-scanning behaviors^2,9^. In previous studies, the head-orientation variability at given location served to identify the cognitive processes underlying spatial decision-making and spatial representation. In (^2^), head-orientation variability served to identify VTE behaviors, whereas in (^9^), it served to identify head-scanning behaviors. However, head-orientation variability is a continuous measure rather than simply a binary marker for behavioral events. We hypothesized that head-orientation variability would be modulated with learning, as observed for other spatial measures (trajectory length, duration and similarity). As expected, IdPhi scores progressively decrease with increasing trial number, suggesting that head-orientation variability is modulated by spatial learning (fig:4).

Evidence for the relationship between VTE behavior and spatial learning have been previously shown^2,6^. In a review of the VTE behavior in rodents, David Redish showed indications that VTE occurs mostly in early stages of spatial learning^6^. Interestingly, quantification of VTE is measured at specific spatial decision points of the environment, whereas navigational performance is based on spatial features over the entire navigational path. Because IdPhi was measured, in our experimental setup, at the very first decision-making intersection, we aimed to understand whether such measure was prognostic of the participants performance of each trajectory. We observed a high correlation between IdPhi at the first intersection and the length and duration of the corresponding trajectory, suggesting that, indeed, occurrence of head-scanning is a predictor of spatial learning.

We sought to understand whether humans, like rodents, benefit from the variability of head-orientation during spatial decision-making, and how it is further related with spatial learning. Furthermore, we devised a virtual environment and experimental protocol where place and response strategies could be used to solve navigation. Our results suggest that humans are capable of adapting their navigational strategy depending on their understanding of the task contingency. Furthermore, we have shown that learning is tightly related with the amount of head-orientation variability, suggesting the functional role of VTE and head-scanning in spatial-representation and navigational knowledge. Together, our results highlight the transferability of results found in the rodent model to the human model when it comes to the behavioral aspects of navigation.

## Acknowledgements

The research leading to these results has received funding from the European Research Council under the European Union’s Seventh Framework Programme (FP7/2007–2013)/ERC grant agreement no (341196) cDAC. We thank Sock Ching Low and Daniel Pacheco for their comments on early versions of the manuscript.

## Author contributions statement

DSP and PFMJV designed the experiment, interpreted the results and wrote the manuscript. DSP implemented the virtual reality setup, acquired the data and analyzed the data.

## Additional information

The authors declare no competing financial interests.

